# Y-profile evidence: close paternal relatives and mixtures

**DOI:** 10.1101/373423

**Authors:** Mikkel M Andersen, David J Balding

## Abstract

We recently introduced a new approach to the evaluation of weight of evidence (WoE) for Y-chromosome profiles. Rather than attempting to calculate match probabilities, which is particularly problematic for modern Y-profiles with high mutation rates, we proposed using simulation to describe the distribution of the number of males in the population with a matching Y-profile, both the unconditional distribution and conditional on a database frequency of the profile. Here we further validate the new approach by showing that our results are robust to assumptions about the allelic ladder and the founder haplotypes, and we extend the approach in two important directions. Firstly, forensic databases are not the only source of background data relevant to the evaluation of Y-profile evidence: in many cases the Y-profiles of one or more relatives of the accused are also available. To date it has been unclear how to use this additional information, but in our simulation-based approach its effect is readily incorporated. We describe this approach and illustrate how the WoE that a man was the source of an observed Y-profile changes when the Y-profiles of some of his male-line relatives are also available. Secondly, we extend our new approach to mixtures of Y-profiles from two or more males. Surprisingly, our simulation-based approach reveals that observing a 2-male mixture that includes an alleged contributor’s profile is almost as strong evidence as observing a matching single-contributor evidence sample, and even 3-male and 4-male mixtures are only slightly weaker.

## Introduction

In [1], we presented a radically simple new approach to the evaluation of weight of evidence (WoE) for Y-chromosome profiles. We showed using simulation that sets of males with the same Y-profile typically number up to a few tens, and rarely more than a few hundreds, almost all of them related within a few tens of meioses. Our simulation model is implemented in open-source and easy-to-use R software malan [2], allowing these distributions to be approximated under different assumptions about the variance in reproductive success (VRS) and the population size and growth rate. We also showed how the distribution of |Ω|, the number of males with the same Y-profile as an alleged source Q, is affected by conditioning on a database count of the profile. In particular, we noted that a zero count in a database of up to a few thousand profiles conveys little information, since from the mutation rate we expect any profile to be rare, which is reflected in the unconditional distribution of |Ω|.

In some cases the Y-profiles of one or more male-line relatives of Q may also be available. This information also affects the distribution of |Ω|, and here we use a simple modification of our simulation model to investigate its effect on the WoE. Any patrilineal relative observed to have a Y-profile not matching that of Q decreases |Ω| in distribution, and hence tends to increase the WoE for Q to be the source of the evidence profile. Conversely a matching relative tends to increase |Ω| and so weaken the WoE. Note that if the relative’s Y-profile differs from that of Q at multiple loci then the proposed biological relationship may be called into question; we do not consider further here the possibility of a mis-specified relationship.

Suppose that, rather than observing a profile matching that of Q, we observe a mixture of the Y-profiles of two or more males such that the profile of Q is “included in the mixture” (every allele in the profile of Q is observed in the mixed profile). Then, because there can be millions of distinct profiles that are included in the mixture, it is typically assumed that the WoE for Q to be a contributor is correspondingly weaker than in a single-contributor case. We show that this intuition is incorrect. This is because the number of distinct Y-profiles that actually arise in a real human population is only a minuscule fraction of the possible profiles given the alleles at each locus. Therefore, although there are many alternative profile combinations that could explain the observed mixture, the great majority of these combinations do not exist in the population, whereas the profile of Q has been observed and is likely to also exist in his close relatives. We show that a 2-male mixture that includes Q has almost exactly the same evidential value as a single-contributor match, and 3-male and 4-male mixtures are only slightly weaker.

Before tackling the above two major goals of this paper, we provide further support for our simulation-based approach by showing that our results are robust to assumptions about the allelic ladder of the mutation model, and the method of allocation of haplotypes to founders. In [1] we assumed an unbounded allelic ladder and that all founders were assigned the same haplotype. Here we adopt more realistic assumptions, but first confirm that this change makes little difference to the results.

## Methods and materials

### Profiling kits, allelic ladders and founder haplotypes

We consider two Y-chromosome short tandem repeat (STR) profiling kits: PowerPlex Y23 (23 loci) and Yfiler Plus (27 loci). As in [1], we continue to consider only integer alleles in our simulations, but they are now bounded by *L* (lower) and *U* (upper). An *L* allele can only mutate to *L+*1, while a *U* allele can only mutate to *U*−1. All other alleles remain equally likely to increase or decrease at a mutation, and the mutation rate is the same for all alleles at a locus. The values of *L* and *U* are specified at each locus corresponding to the integer alleles in YHRD.org release 55 [3] (see Fig. 1). For comparison, we also considered a tiny ladder of size 3 (alleles −1, 0 and 1).

**Figure 1:**
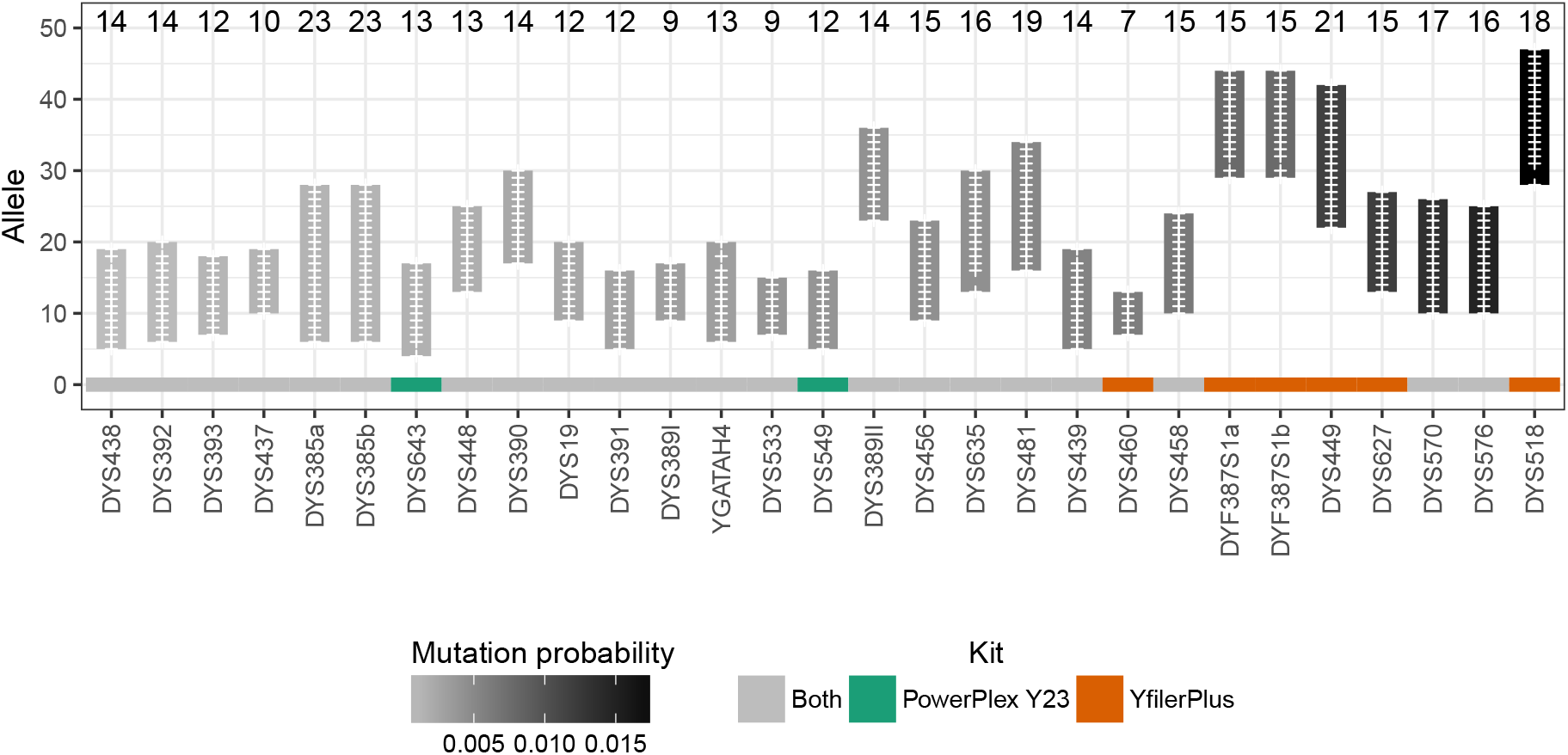
Profiling kits and allelic ladders. Only integer alleles are included, not alleles with partial repeats. Vertical bars indicate the ladders, with a “+” for each observed allele, shaded according to the estimated locus mutation rate per generation as indicated in the legend. The size of each allelic ladder is given above the bar. Data are from YHRD.org release 55 [3].

In [1], all founders in the population simulation got the same haplotype. This implied that if few mutations occurred since their founders, two live individuals could have matching haplotypes despite descending from distinct founders. We set the number of generations such that this was very unlikely, but for further realism we consider here two different ways of assigning haplotypes to founders:

- Uniformly random choices from the (integer) allelic ladder, independently at each locus.
- Haplotypes sampled at random with replacement from a contemporary Danish database of 185 males [4]. We removed profiles with three alleles at DYF387S1, non-integer alleles, or null alleles, leaving 181 PowerPlex Y23 profiles and 171 Yfiler Plus profiles.

### Population simulations

We used our R package malan [1, 2, 5] to simulate 10 population genealogies: an initial population of 5,000 Y chromosomes reproduces for 100 generations, followed by growth at a rate of 2% per generation for 150 generations, creating a final population size of 102K. Thus, the number of live males (total of final three generations) is close to 300K. The VRS was fixed here at 0.2; see Fig. A1 for the distributions of the number of sons and brothers of each male. To each population simulation we applied two allelic ladders (bounded/unbounded) for each of three assignments of founder haplotypes (same/random/database) and each of two kits (PowerPlex Y23/Yfiler Plus). The mutation process was replicated 10 times. Following [1], we used mutation count data [3] with a Beta(1.5, 200) prior distribution at each locus to obtain a posterior distribution from which the mutation rate was sampled, independently over loci.

In each simulation 5,000 males (Q) were drawn at random and for each we recorded |Ω|, the number of live males with the same haplotype (including Q). Thus, for each of the 12 ladder / founder / kit combinations, the distribution of |Ω| was estimated based on 10 (genealogies) × 10 (mutation replicates) × 5, 000 (choices of Q) = 5 × 10^5^ cases. In each simulation, information about the profiles of close paternal relatives of Q was also recorded, so that we could approximate the distribution of |Ω| conditional on the profile status of different relatives.

For comparison, we include below results from [1] which used a slightly different population simulation that we now briefly recap: 250 generations; growth of 2% in all generations; initial population size of 7,365 rising to 10^6^ in the final generation (in our new simulations, the growth rate is the same but for fewer generations, and initial and final population sizes are both smaller). Ten genealogies were simulated; mutation rates were sampled 100 times per genealogy (c.f. 10 here); only an unbounded allelic ladder was considered with the same haplotype for each founder, and 1,000 Q were sampled per simulation.

### Mixed profiles

In general, the preferred measure of the WoE for Q to be a contributor to an evidence sample is the likelihood ratio (LR) [6]. When the evidence sample shows exactly his profile *q*, the LR is the inverse of a (conditional) match probability, but if we know the Y-haplotype counts in the population of *N* alternative sources of the evidence profile, then the conditioning is irrelevant and the LR simplifies to

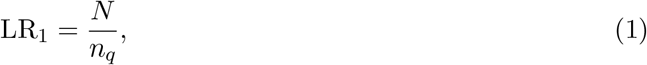

where we introduce the notation *n_a_* for the count of haplotype *a* in the population. In [1], we did not recommend reporting LR_1_, because the population size relevant to a crime scenario is often highly uncertain. Instead we recommended reporting an estimate of the haplotype count.

Suppose now that the evidence profile *m* has two different alleles at *h* loci, and no more than two alleles at any locus. There are 2^*h*−1^ possible profile pairs that could have produced the mixture, which is the number of ways of choosing one allele from *m* at each locus, and ignoring the order of the resulting profile pair. Suppose also that an alleged contributor Q has profile *q* that is included in *m*. Then a relevant LR to consider compares the hypothesis *H_p_*, that *m* arises from Q and an unknown male U, relative to the alternative *H_d_* that *m* arises from two unknown males [6]. Under *H_p_* the profile *u* of *U* can be inferred from *q* and *m*, without error if we assume no missing data or null alleles, and no duplications or heteroplasmy, so both males have exactly one allele at each locus. Still assuming that the *n_a_* are known in the population of possible sources of *m*, we have:

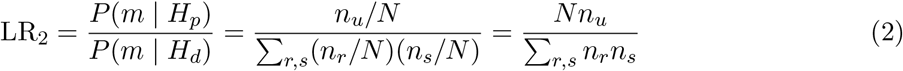

where the summation is over the 2^*h−*1^ unordered pairs of profiles (*r, s*) that combine to give *m*. LR_2_ can be interpreted as the probability that two profiles drawn at random in the population form *m*, divided by the probability that a single profile drawn at random forms *m* when combined with *q*. If *n_q_*, *n_r_* and *n_s_* are all of comparable magnitude then LR_2_ *≈* LR_1_/2^*h−*1^. For current Y-profiles, 2^*h−*1^ can exceed one million, and so the WoE from a mixed evidence profile is usually considered to be much weaker than from a single-contributor evidence profile.

[3, 7] compute (2) directly, using observed database fractions in place of population fractions of the form *n_a_/N*. However, databases are not large enough for accurate estimation of these, small, fractions. More importantly, the relatedness of males with the same haplotype means that they may be clustered geographically and socially, meaning that the available databases are unlikely to accurately represent the population of possible sources of the evidence profile in a specific case.

[8] used [9, 10] to obtain improved estimates of population fractions by modelling the haplotype distribution as composed of clades of haplotypes each of which has arisen from one ancestral haplotype by a small number of single-step mutations. Within each clade, independence is assumed across loci and haplotype probabilities are computed using a mixture of discrete Laplace distributions. The population fraction of the haplotype is obtained as a weighted sum over the clades (the weights correspond to the prior probability that a haplotype in the population originates from that clade).

[11] further develop the clade idea, but recognise the importance of the fact that profile *q* has been observed, which is typically not the case for other profiles included in the mixture. They introduce a “haplotype centred” method to compute the LR, which uses the insight that, given the observed profile of Q, the most likely source of a matching or similar profile is in a close patrilineal relative of Q, as previously noted by [12].

The approach proposed here is different but based on a similar insight. We note that although (*q, u*) is just one among many profile pairs that could contribute to the summation in (2), if *q* is the only reference profile available to the investigation that is included in *m*, then (*q, u*) is expected to provide the largest contribution to the sum. The number of different Y-profiles that actually arise in any human population is a tiny fraction of the profiles that are possible. For example, just the integer alleles of the 27 Yfiler Plus loci shown in Fig. 1 can generate more than 10^31^ distinct profiles, whereas the worldwide human population is < 10^10^. Thus, a random possible profile is extremely unlikely to actually exist. In contrast, the fact that profile *q* has been observed in Q implies that we expect it to exist in multiple male-line relatives of Q. Although we have no *a priori* evidence for the existence of profile *u*, it is much more likely that one unobserved profile exists in the population than that two unobserved profiles *r* and *s* both exist.

To quantify the extent to which the profile pair (*q, u*) dominates the summation in (2), we use the 500K malan simulations described above in the case of bounded allelic ladder and database founder haplotypes. From each simulation, we sample pairs of live males Q and U and form the mixed profile *m*. We then search the live population for other pairs of males whose mixed profile is also *m*. As for the single-contributor case, because of the problem of specifying *N* in practice, instead of LR_2_ we recommend reporting

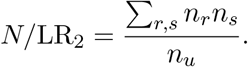

Similarly we search for triples of males with mixed profile *m* matching that from Q and two other randomly-selected males, and report

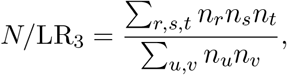

where each (*r, s, t*) in the sum is a triple of profiles that combine to form *m*, and (*u, v*) is a pair of profiles that when combined with *q* form *m*. The expression for *N/*LR_4_ is analogous.

## Results

### Robustness to allelic ladder and founder haplotypes

Quantiles of the distribution of |Ω|, the number of males with the same Y-profile as Q, are shown in Table 1 for the different allelic ladders and methods of assigning founder haplotypes. See Fig. A2 for plots. The distributions are similar for all conditions considered here. The biggest, but still small, impact arises from using a tiny ladder of size three at each locus.

**Table 1:** Estimated quantiles of the distribution of |Ω|, the number of males with the same Y-profile as Q. See text for explanation of the allelic ladders (column 1) and the methods of assigning founder haplotypes (column 2). Row 3 gives results previously published in [1], similar to the case of Row 2 but with a slightly different demographic model as discussed in the text. PP23 = PowerPlex Y23; YP = Yfiler Plus.

| Allelic<br>ladder | Founder<br>haplotypes | 95% quantile |  | 99% quantile |  |
| --- | --- | --- | --- | --- | --- |
|  |  | PP23 | YP | PP23 | YP |
| Unbounded | Random | 73 | 41 | 114 | 64 |
| Unbounded | Same | 73 | 41 | 114 | 64 |
| Previously published |  | 73 | 41 | 115 | 63 |
| Bounded | Random | 73 | 41 | 115 | 64 |
| Bounded | Database | 73 | 41 | 115 | 64 |
| Bounded | Same | 74 | 41 | 115 | 63 |
| $\{-1, 0, 1\}$ | Random | 77 | 42 | 120 | 65 |

### Profiled male-line relatives

If either the father or the paternal grandfather is observed not to match Q, then the distribution of |Ω| gives greatly increased support to low values, whereas a match shifts the distribution slightly towards higher values compared with the unconditional case (Fig. 2 and Table 2).

**Figure 2:**
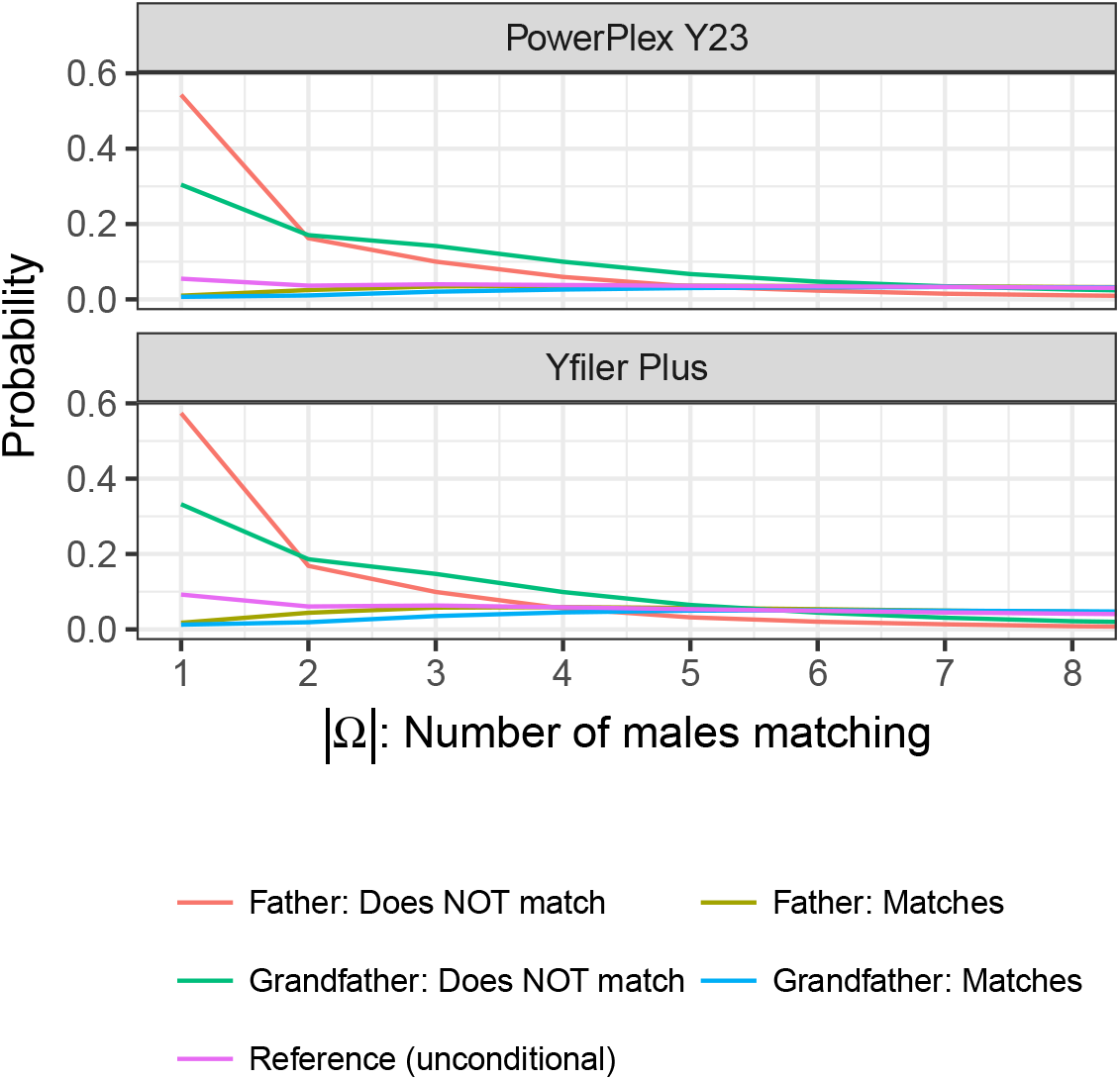
Distribution of |Ω| given father/grandfather profile match information. “Unconditional” means without information about the Y-profile of any relative of Q. The other lines correspond to match/mismatch information as indicated in the legend.

**Table 2:** Estimated quantiles of the distribution of |Ω| given match information about specified patrilineal relatives. See Figs 2, 3 for plots. PP23 = PowerPlex Y23; YP = Yfiler Plus.

| Data | 95% quantile |  | 99% quantile |  |
| --- | --- | --- | --- | --- |
|  | PP23 | YP | PP23 | YP |
| Father: Does NOT match | 9 | 7 | 30 | 15 |
| Grandfather: Does NOT match | 13 | 10 | 32 | 18 |
| Random uncle: Does NOT match | 48 | 27 | 90 | 49 |
| Random brother: Does NOT match | 56 | 32 | 97 | 54 |
| Random cousin: Does NOT match | 58 | 33 | 98 | 55 |
| Random son: Does NOT match | 72 | 40 | 113 | 61 |
| Reference (unconditional) | 73 | 41 | 115 | 63 |
| Random son: Matches | 74 | 42 | 115 | 65 |
| Father: Matches | 76 | 43 | 117 | 65 |
| Random brother: Matches | 77 | 44 | 119 | 67 |
| Grandfather: Matches | 78 | 45 | 120 | 68 |
| Random uncle: Matches | 80 | 47 | 122 | 70 |
| Random cousin: Matches | 81 | 48 | 123 | 71 |

Fig. 3 and Table 2 describe the distribution of |Ω| given information about the match status of a specified patrilineal relative, or that there is no relative of the specified type. The effect of the latter information is seen to be intermediate between match and mismatch for that relative.

**Figure 3:**
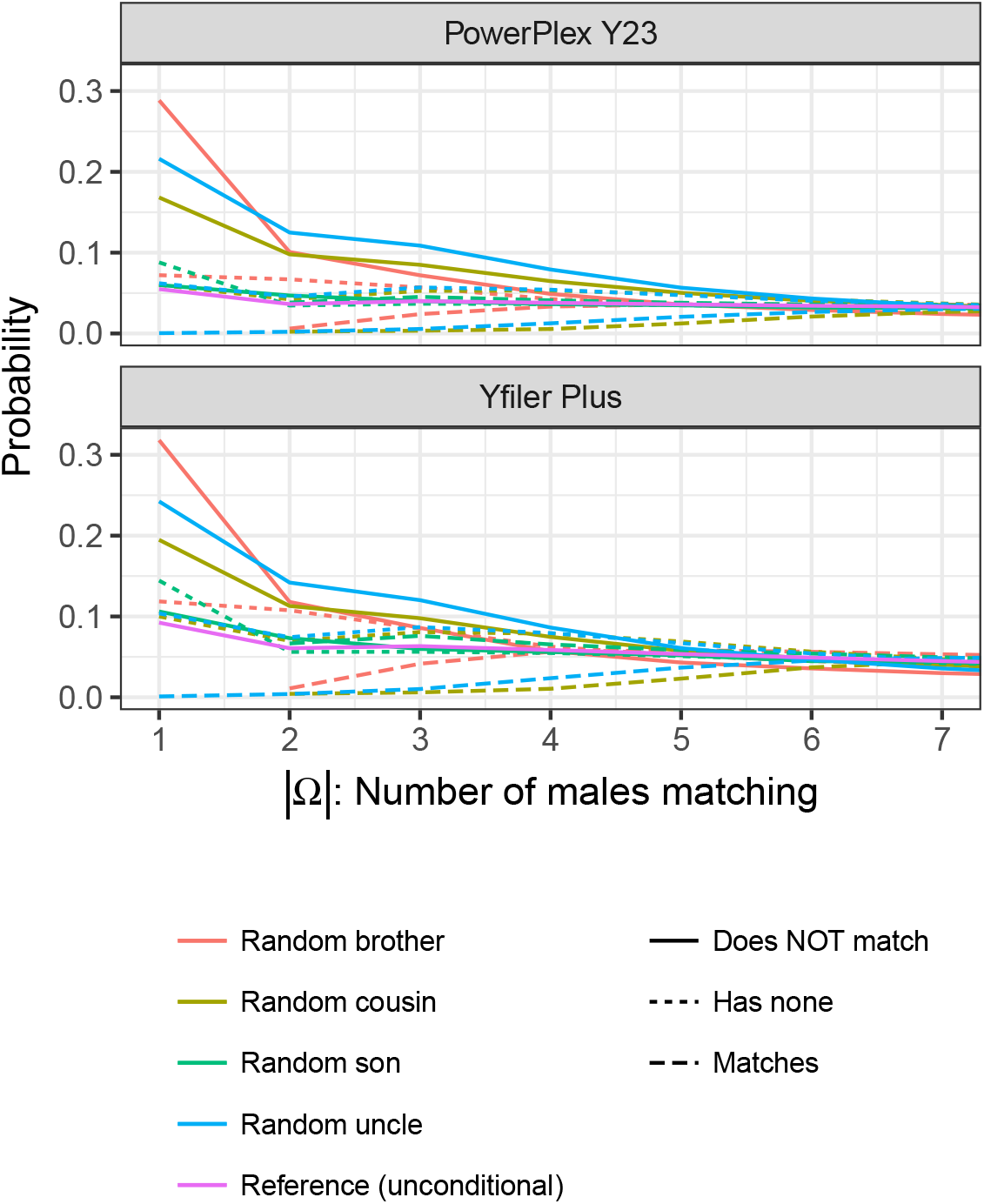
Distribution of |Ω| given that Q has no son, brother, patri-lineal cousin or paternal uncle, or match information from a random one of them. Each curve is coded by its colour and line type (see legend); no information is available about male-line relatives other than the one specified. The reference line corresponds to no information from relatives of Q.

While broadly in line with intuition, these results hold some surprises. In general, the closer the relative the greater is the effect of a mismatch in decreasing the distribution of |Ω|, but the direction in time of the relationship is also important: a mismatching father is more important than a mismatching son, because the father has more descendants. Similarly, a grandfather is more informative than a brother. Moreover, a mismatching brother has only slightly greater impact on the quantiles of |Ω| than a mismatching cousin: a brother relationship is closer but a cousin relationship traverses one extra generation backward in time and is informative about the grandfather. For matches, more distant relatives are more informative, with a matching cousin being most informative among the relationships considered here, but because a cousin is expected to match the impact of this information on the distribution of |Ω| is modest.

In Fig. 4 the match/mismatch information comes from all the brothers of Q, for Q with between one and three brothers. For Q with two or three brothers, all of them with a Y-profile different from Q, the distribution of |Ω| is similar to the case that Q is found not to match his father.

**Figure 4:**
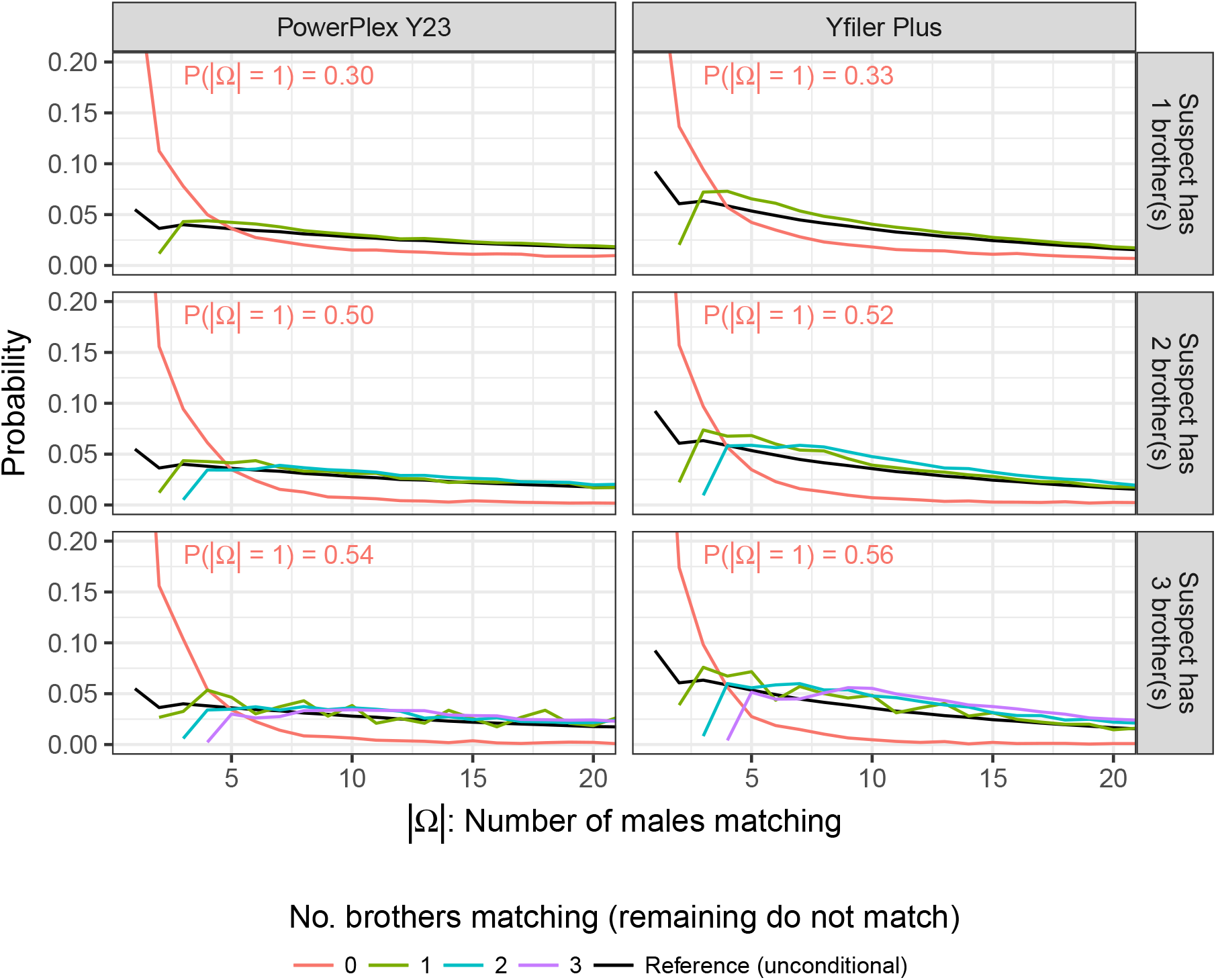
Distribution of |Ω| given match information for all brothers when Q has up to three brothers. The value of *P* (|Ω|=1) when all brothers mismatch is off the plot and is given numerically.

### Mixed profiles

For 97% of 2-male Yfiler Plus (YP) mixed profiles, the mixture cannot be formed from any other pair of profiles that actually exists in the population (Table 3, second row). In that case, a mixed evidence profile is equivalent to an evidence profile exactly matching *q*. This equivalence holds for 93% of 2-male PowerPlex Y23 (PP23) mixtures; for 56% (YP) and 42% (PP23) of 3-male mixtures and for 20% (YP) and 10% (PP23) of 4-male mixtures, respectively.

**Table 3:**
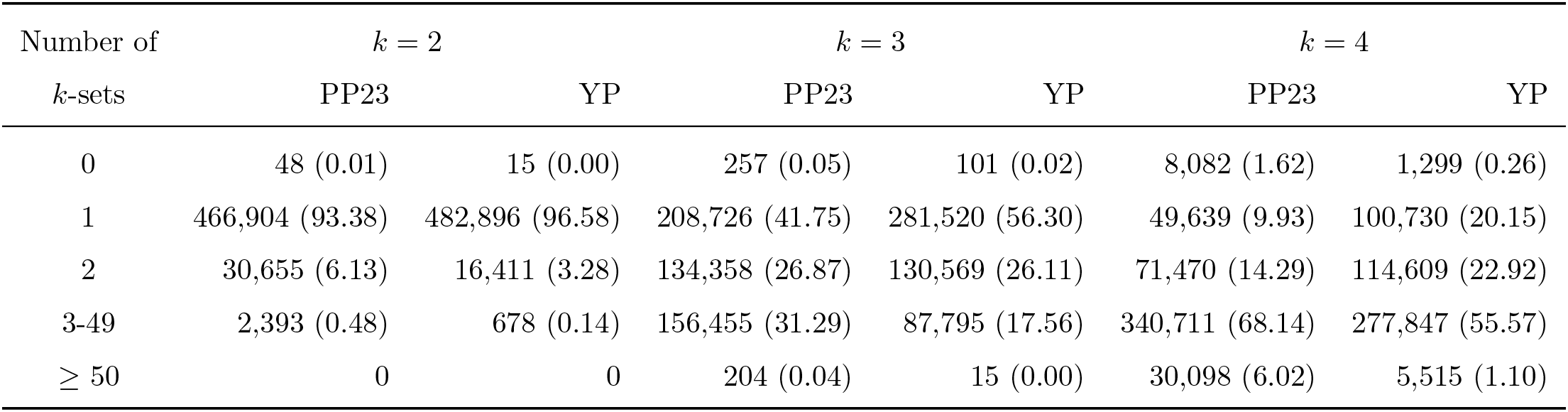
Distribution of the number of distinct *k*-sets of profiles yielding a given *k*-male mixed Y profile. Each cell records the count (%) of 500K simulated *k*-male mixtures that could be obtained in the number of different ways indicated in the first column. The first row (0) corresponds to when the mixture is not recognised as a *k*-male mixture because no locus had *k* alleles. For *k* = 2 this only happens when the two contributors have the same profile. The second row (1) corresponds to cases when the profiles generating the mixture form the only *k*-set of profiles in the live population that combine to form that mixture. PP23 = PowerPlex Y23; YP = Yfiler Plus.

| Number of<br>$k$-sets | $k = 2$ | | $k = 3$ | | $k = 4$ | |
| --- | --- | --- | --- | --- | --- | --- |
|  | PP23 | YP | PP23 | YP | PP23 | YP |
| 0 | 48 (0.01) | 15 (0.00) | 257 (0.05) | 101 (0.02) | 8,082 (1.62) | 1,299 (0.26) |
| 1 | 466,904 (93.38) | 482,896 (96.58) | 208,726 (41.75) | 281,520 (56.30) | 49,639 (9.93) | 100,730 (20.15) |
| 2 | 30,655 (6.13) | 16,411 (3.28) | 134,358 (26.87) | 130,569 (26.11) | 71,470 (14.29) | 114,609 (22.92) |
| 3-49 | 2,393 (0.48) | 678 (0.14) | 156,455 (31.29) | 87,795 (17.56) | 340,711 (68.14) | 277,847 (55.57) |
| $\geq 50$ | 0 | 0 | 204 (0.04) | 15 (0.00) | 30,098 (6.02) | 5,515 (1.10) |

**Table 4:** Estimated quantiles of the distribution of *N*/LR*_k_* for *k* = 1,…, 4. See Fig. 5 for plots. PP23 = PowerPlex Y23; YP = Yfiler Plus.

|  | 95% quantile |  | 99% quantile |  |
| --- | --- | --- | --- | --- |
|  | PP23 | YP | PP23 | YP |
| $N/\text{LR}_1$ | 72 | 40 | 113 | 63 |
| $N/\text{LR}_2$ | 72 | 41 | 119 | 64 |
| $N/\text{LR}_3$ | 82 | 45 | 139 | 71 |
| $N/\text{LR}_4$ | 99 | 51 | 177 | 83 |

To explain this startling result in simpler terms, imagine a mixture in which alleles 1 and 2 are observed at each of 25 loci and an alleged contributor Q has allele 1 at every locus. Then the number of possible distinct profile pairs contributing to the mixture is 2^24^ or almost 17 million. However, under our simulation model, it is highly probable that the two profiles contributing to the mixture are (1, 1,…, 1) and (2, 2,…, 2): the other 17 million possible profile pairs are collectively unlikely to exist in the population.

Table 3 does not answer the WoE problem for mixed evidence profiles, but it helps explain Fig. 5 which shows the distribution in our simulations of *N*/LR_*k*_ for *k* = 1,…, 4 (ignoring those counted in the first row of Table 3). As expected, the distribution is shifted towards higher values as *k* increases, reflecting reduced WoE as the number of contributors to the evidence sample increases. What is striking and counter-intuitive is that the reduction in WoE is so slight. One guide to the correct intuition is that, for example when *k* = 4, if there are many quadruples of males in the population whose profiles combine to make *m*, then there are also many triples that when combined with *q* also make *m*: the Spearman correlation between the number of quadruples and the number of triples is around 0.85 for both kits.

**Figure 5:**
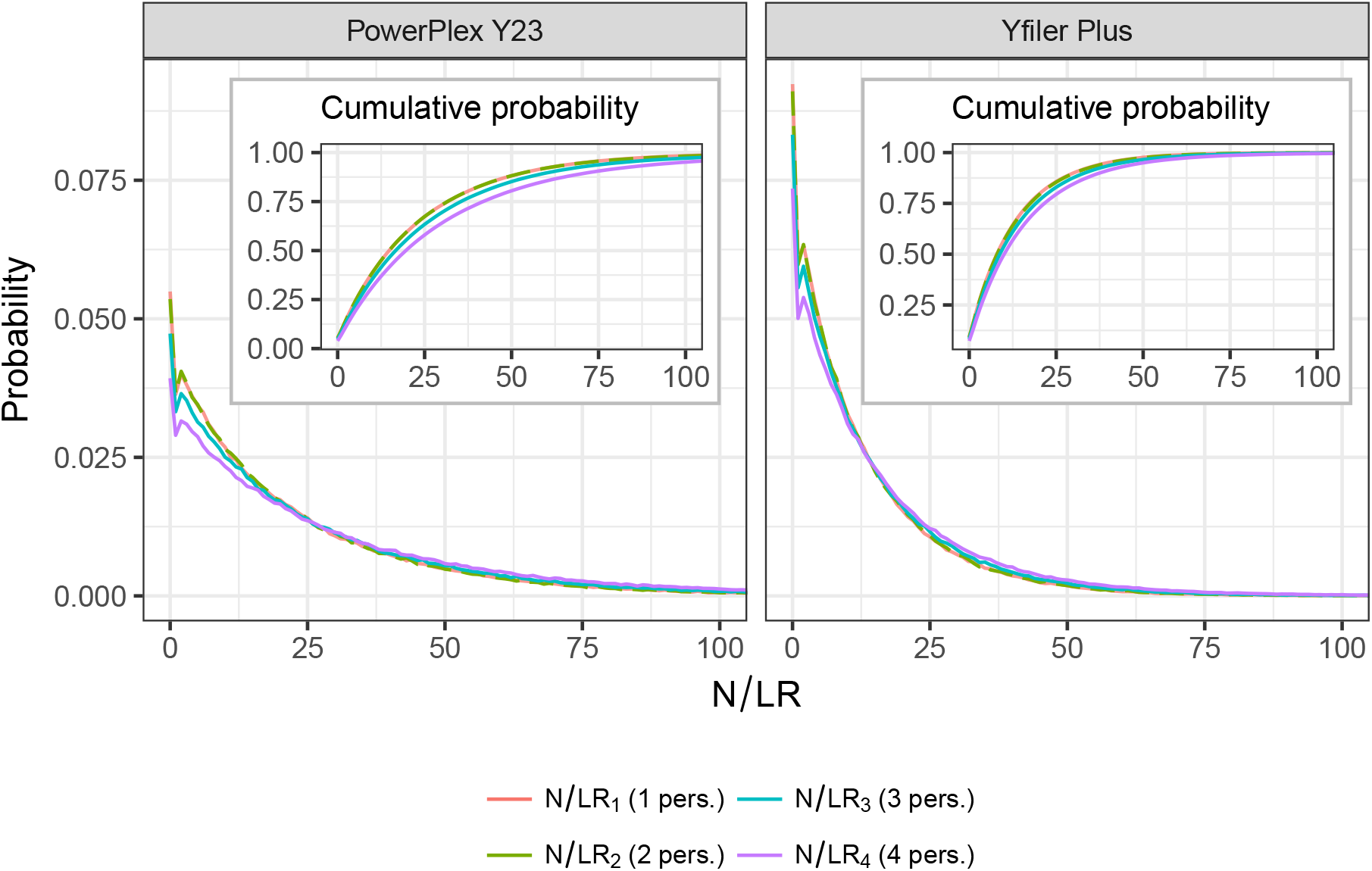
The distribution of *N/*LR_*k*_ for *k* = 1,…, 4. The case *k* = 1 corresponds to Fig. 4 of [1], and describes the distribution of the number of males with Y-profile matching that of a single-contributor evidence profile. The other curves describe the distribution of an analogous measure of WoE (see methods) when a reference profile *q* is included in a *k*-male mixture, *k* = 2, 3, 4. The red and green curves (*k* = 1, 2) are almost indistinguishable and are shown with alternating colours. See Table 4 for key quantiles.

## Discussion

We have further developed our new and powerful simulation-based approach to assessing the weight of Y-profile evidence [1]. We have extended it to allow conditioning on the profiles of some male-line relatives of the alleged contributor Q, and to evidence samples that include DNA from up to four males. The simulations underlying our results can be performed for any profiling kit, which is demonstrated in a vignette in the R package malan [2].

The results for conditioning on male-line relatives broadly match intuition though with some surprising aspects. If either the father or grandfather of Q is observed to have a Y-profile different from Q, then the number of matching males |Ω| is greatly reduced, and consequently the Y-profile evidence is strengthened in favour of Q being the source (Fig. 2). More generally, the distribution of |Ω| is reduced for any observed mismatch with a male-line relative of Q. The converse is also true: observed relative matches reduce the WoE. However, the magnitude of effect is not symmetric. From Table 2, we see that the observation of a mismatching father has the greatest impact on the distribution of |Ω|, whereas a matching father has little impact.

Our most striking result is that the observation that the profile of Q is included in a mixed evidence profile with up to four contributors is almost as strong evidence for Q to be a contributor as is a match with a single-contributor evidence profile. In particular, being included in a 2-male mixture has virtually the same evidence value as a single-contributor match. This property was not apparent using previous approaches to evaluating mixtures. Although there are many other sets of profiles that could generate the mixture, if a possible profile has not been observed it is very unlikely to actually exist in the population. It follows that a 2-male mixed evidence profile can be presented to a court in terms of an equivalent single-contributor profile, using the suggestions we made in [1]. A similar approach may also be feasible for 3-male and 4-male mixtures.

The results reported here have been obtained using one model, but malan can be used to investigate alternative mutation models and demographic scenarios. As we noted in [1], almost all Y-profile matches are between males who are related to within a few tens of meioses. It follows that our results are robust to the mutation mechanism, with only the mutation rate being important. Moreover the number of matching males is typically up to a few tens, which is small relative to the population size and so our results are also reasonably robust to details of the demographic model [1]. We have confirmed here that the distribution of |Ω| is robust to assumptions about the allelic ladder and the allocation of founder haplotypes.

Overall, these results further advance the case for use of our new simulation-based paradigm for Y-profile evidence, introduced in [1]. We have demonstrated here that our approach is flexible enough to incorporate new kinds of evidence, and it leads to important new insights about the strength of Y-profile evidence.

## Acknowledgements

This work was supported in part by the Otto Mønsted Foundation and a short term fellowship from the International Society for Forensic Genetics (ISFG).

## Supplementary information

**Figure A1:**
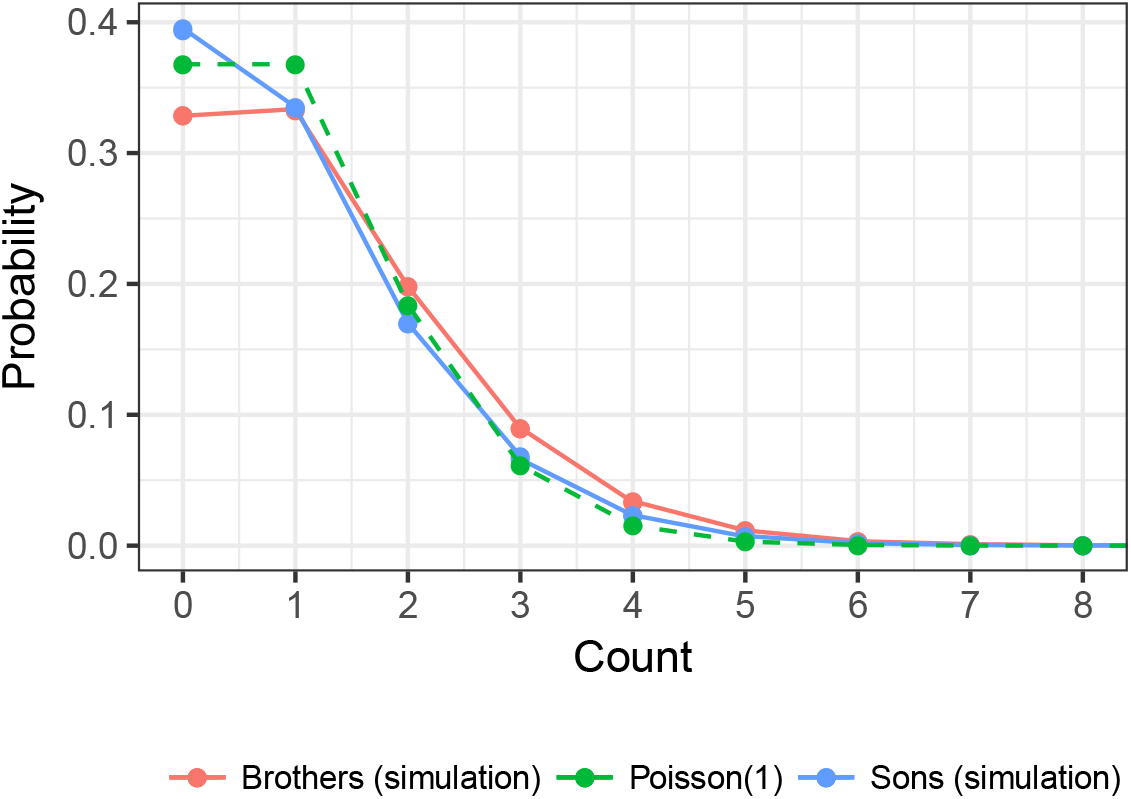
The distribution of the numbers of sons and brothers of each male. Note that these distributions are affected both by the variance in reproductive success (here, VRS = 0.2) and the population growth rate (0.02). The Poisson(1) distribution is plotted for comparison.

**Figure A2:**
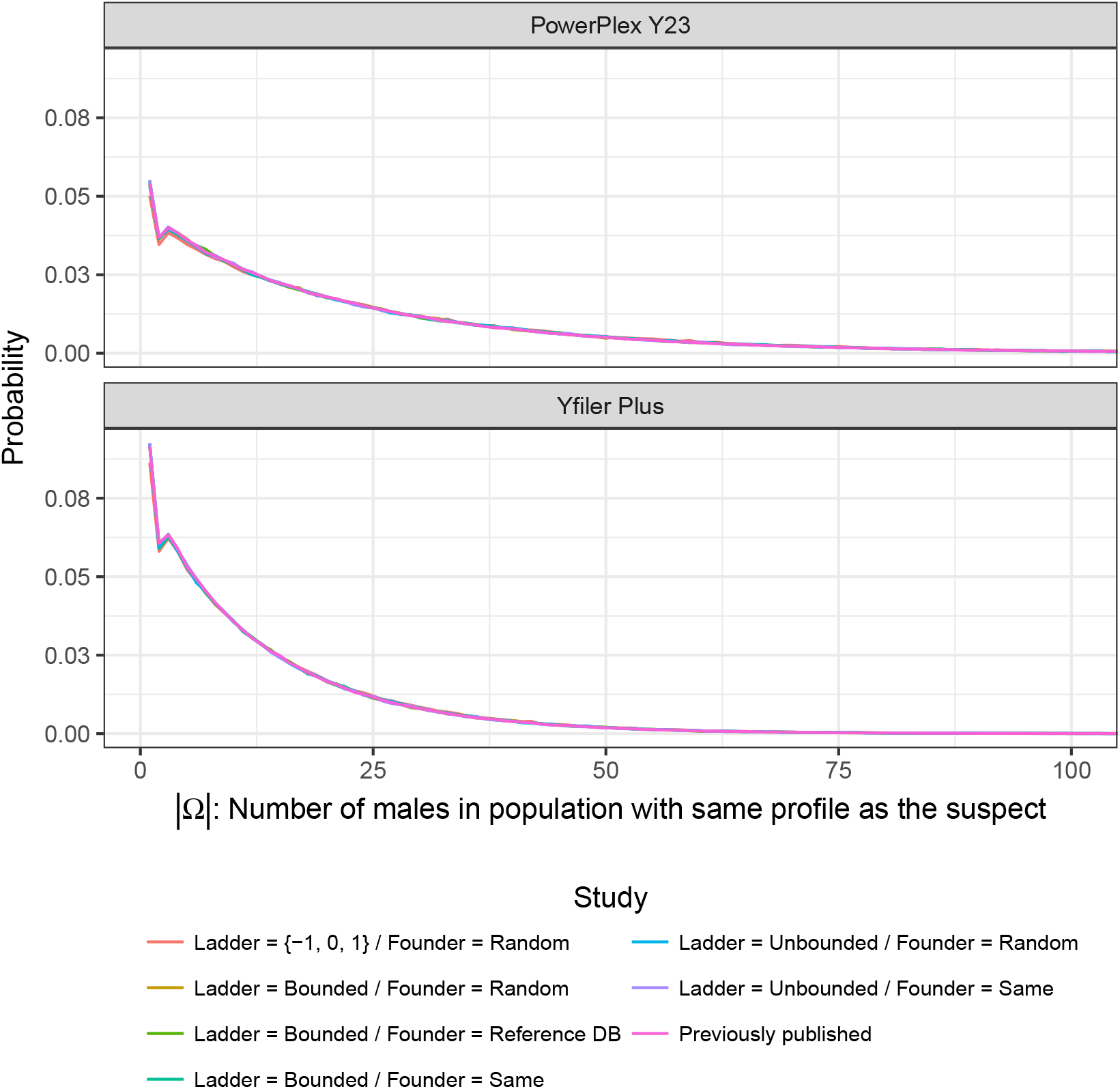
Distributions of |Ω| under six models for allelic ladder and founder haplotype assignment. The distribution reported in [1] is also shown.

